# Unlocking Precision Gene Therapy: Harnessing AAV Tropism with Nanobody Swapping at Capsid Hotspots

**DOI:** 10.1101/2024.03.27.587049

**Authors:** Mareike D. Hoffmann, Joseph P. Gallant, Aaron M. LeBeau, Daniel Schmidt

**Author notes:** To whom correspondence should be addressed: Daniel Schmidt: 420 Washington Ave SE, Minneapolis, MN 55455, USA, Correspondence may also be addressed to: Mareike D. Hoffmann: Washington Ave SE, Minneapolis, MN 55455, USA.

## Abstract

Adeno-associated virus has been remarkably successful in the clinic, but its broad tropism is a practical limitation of precision gene therapy. A promising path to engineer AAV tropism is the addition of binding domains to the AAV capsid that recognize cell surface markers present on a targeted cell type. We have recently identified two previously unexplored capsid regions near the 2-fold valley and 5-fold pore of the AAV capsid that are amenable to insertion of larger protein domains including nanobodies. Here, we demonstrate that these hotspots facilitate AAV tropism switching through simple nanobody replacement without extensive optimization in both VP1 and VP2. We demonstrate highly specific targeting of human cancer cells expressing fibroblast activating protein (FAP). Our data suggest that engineering VP2 is the preferred path for maintaining both virus production yield and infectivity. Our study shows that nanobody swapping at multiple capsid location is a viable strategy for nanobody-directed cell-specific AAV targeting.

## INTRODUCTION

Adeno-associated virus (AAV) is a compact 25 nm virus known for its favorable clinical characteristics, such as low pathogenicity and the ability to induce long-term expression in both dividing and non-dividing cells. These features make it a promising candidate for applications in gene and cell therapy (reviewed in (1). Its single-stranded DNA genome, spanning 4.7 kb, encodes two genes, *rep* and *cap*, flanked by inverted terminal repeats. The *cap* gene produces three viral proteins (VP1, VP2, and VP3) from the same open reading frame. VP2 is a N-terminal truncated version of VP1, and VP3 is a further truncated version of VP2 (2, 3). The capsid, formed by 60 VP monomers, exhibits an icosahedral structure with an average ratio of 1:1:10 for VP1, VP2, and VP3, respectively (4–6).

The capsid’s distinctive features include a protruding, cylindric pore at the 5-fold interface, a valley extending toward the 2-fold interface around the pore, and protrusions at the 3-fold interface, which play a crucial role in mediating target cell receptor binding (7). Despite high conservation in the overall structure and topology across serotypes, structural analyses have identified nine variable regions (VR1-9) on the capsid surface (8). VR4 and VR8 have been particularly targeted in AAV capsid engineering to evade neutralizing antibody binding (9) and re-direct viral tropism (10–15).

While VR8 has been predominantly utilized for peptide insertions, a library screen by Judd et al. (16) revealed that VR4 can accommodate the fluorescent protein mCherry. Subsequent to this discovery, various protein domains with re-targeting capabilities, including DARPins (17), HUH-tags (18), and nanobodies (19, 20), were successfully incorporated into VR4. Nanobodies, originating from camelids and characterized by their small size (15 kDa), specificity, stability, and ability to serve as targeting ligands for chemotherapy drugs, radionuclides, or toxins (21), stand out among these options. Therefore, there is considerable interest in optimizing the incorporation of nanobodies into the AAV capsid to enhance cell type specific AAV targeting.

In previous work, we have had performed a domain insertion library screen by incorporating domains with re-targeting abilities, including a GFP nanobody, in between every two amino acid residues of VP1 protein of AAV-DJ (22). We demonstrated that nanobody insertions are tolerated not only in the tip of the 3-fold protrusion (VR4), but also several positions lining the 2-fold valley as well as the 5-fold interface of an AAV capsid. While insertion of a GFP nanobody into AAV increased infectivity towards cells expressing GFP on their cell surface, it was unclear if this nanobody-mediated re-targeting generalizes to different nanobodies, implying modularity, or whether any optimization is required to achieve high infection specificity.

To address these questions, we selected seven positions near the 2-fold valley and 5-fold interface alongside the benchmark insertion position in VR4. Insertions were made into either the VP1 or VP2 protein of AAV-DJ with two different types of linkers. With an eye toward clinically-relevant AAV re-targeting, we chose a nanobody targeting fibroblast activating protein (FAP), which has emerged as a promising cancer target in recent years (23–25). While most of our FAP nanobody insertion variants have weaker infection efficacy on cells lacking the FAP receptor (i.e., off-target cells), most of the variants (six out of the eight positions tested) surpassed the DJ control in FAP receptor-positive (‘on-target’) cells with a specificity gain of up to 18-fold for the best variant: VP2-N262 with asymmetric linkers. These findings reinforce the feasibility of nanobody-mediated AAV targeting for cell and gene therapy applications.

## MATERIALS AND METHODS

### Cloning

Oligos and gBlocks were obtained from IDT, restriction enzymes and T4 DNA ligase from NEB and PCRs were done using the PrimeSTAR Max DNA polymerase from Takara Bio. PCR products were purified using the DNA Clean & Concentrator-5 Kit (Zymo Research) or the Zymoclean Gel DNA Extraction Kit (Zymo Research) if PCR products were analyzed on 1 %TAE agarose gels. Post cloning, plasmids were transformed into NEB® Stable Competent *E. coli* cells, before plated on LB plates containing carbenicillin at a concentration of 100 μg/ml. Plasmids were isolated using the Zyppy Plasmid Miniprep Kit (Zymo Research), according to the manufacturer’s instructions. All plasmids used in this study are listed in Table S1. For the cloning of the FAP nanobody insertion variants, the BsmBI restriction site was eliminated from the plasmid containing the rep2-capDJ (AAV-DJ) gene sequences by introducing a silent mutation using mutagenesis PCR at position D178 of the cap-DJ gene. The plasmids DJ-VP1 and VP2 were obtained by mutating the start codons for VP2/3 (T138A, M203K, M211L, M235L) and VP1/3 (M1K, M203K, M211L, M235L), respectively. The for the AAV productions obligatory plasmids complementing the missing VPs, DJ-VP2/3 and DJ-VP1/3, were cloned similarly by mutating the start codons for VP1 (M1K) and VP2 (T138A), respectively. FAP or GFP nanobody insertion plasmids were generated using golden gate assembly (26) and the BsmBI restriction enzyme. FAP (Figure S5) and GFP (27) nanobody sequences were human codon-optimized and ordered as gBlocks with either symmetric or asymmetric linkers. Nanobody and linker sequences are given in Figure S1. Post cloning, sequences were verified by Plasmid-EZ sequencing (Azenta Life Sciences).

### Tissue culture

The cell lines CWR-R1-enzalutamide resistant/luciferase^+^ (stably expressing a firefly luciferase), and CWR-R1-enzalutamide resistant/luciferase plus FAP (additionally expressing a human FAP receptor) were provided by the LeBeau lab from the University of Wisconsin (28). Both cell lines and 293AAV cells (Cell Biolabs) were cultured in DMEM (Gibco) supplemented with 10 % fetal bovine serum (Gibco), 4.5 g/L D-glucose, L-glutamine, 110 mg/L sodium pyruvate, and 100 U per mL penicillin/100 μg per mL streptomycin (Gibco).

Media for R1-FAP cells was additionally supplemented with 3 μg/mL puromycin (ApexBio Technology). Cells were kept in a humidified incubator at 37 °C and 5 % CO_2_ and passaged every 2-4 days when reaching a confluency of 70-80 %.

### AAV crude lysate production

293AAV cells were seeded into 6-well plates at a density of 500,000 cells per well. 24 hours later, cells were transfected with 2.5 μg DNA using PEI and an equimolar ratio of the plasmids necessary for the respective AAV production: (i) an Adenohelper plasmid, (ii) a nanoLuciferase encoding plasmid, (iii) a plasmid encoding rep2 and capDJ with start codons of either only VP1 or only VP2, but with a FAP-nanobody insertion, and (iv) a plasmid encoding rep2 and capDJ encoding the VP proteins needed for complementation. Three days post transfection, cells were harvested by flushing off the cells by pipetting and spun down for 5 min at 400 xg. Cells were resuspended in PBS (pH 7.4, Gibco) and then subjected to five freeze and thaw cycles by alternating between liquid nitrogen and a 37 °C water bath. Cell debris was pelleted by centrifugation at 17,000 xg at 4 °C for 10 min. The supernatant containing the AAV particles was stored at -20 °C until use.

### qPCR

Production titers of crude lysate samples were determined as follows. 2 μl of the crude lysates were mixed with PBS, supplemented with 2mM MgCl_2_, and 0.1 μl ultrapure Benzonase Nuclease (Sigma-Aldrich) was added. Samples were incubated at 37 °C for 30 min to digest DNA that was not protected by AAV capsids. Next, 5 μl 10x Proteinase K buffer (100 nM Tris-HCl, pH 8.0, 10 mM EDTA, and 10 % SDS) and 1 μl Proteinase K (20 mg/ml; Zymo Research) were added inhibiting the Benzonase and digesting proteins including the AAV capsid to free the ssDNA. Afterwards, samples incubated for 20 min at 50 °C, followed by heat inactivation of the Proteinase K for 5 min at 95 °C. The viral ssDNA was purified using the DNA Clean & Concentrator-5 Kit (Zymo Research) according to the manufacturer’s instructions for ssDNA purification. All samples were diluted 1:500 in H_2_O prior to qPCR, which was run using a QuantStudio5 Real-Time PCR System (Applied Biosystems) and by using the PowerUp SYBR Green Master Mix (Applied Biosystems), following the manufacturer’s instructions. A primer set binding within the CMV-enhancer (forward: AACGCCAATAGGGACTTTCC, reverse: GGGCGTACTTGGCATATGAT; (29) of the transgene expression cassette was used. To calculate the viral titer in vg/ml, a plasmid standard at a known concentration also containing a CMV-enhancer was used.

### Luciferase assay

R1 and R1-FAP cells were seeded into 96-well plates at a density of 12,500 cells per well, while SK-MEL-24 cells were plated at a density of 10,000 cells per well. The next day, the media was replaced, and cells were transduced at the indicated multiplicity of infection (MOI) with AAV crude lysates. 48h post transduction, the media was aspirated, cells washed with 100 μl PBS (pH 7.4, Gibco) per well and then 25 μl PBS and 25 μl of the Nano-Glo® Luciferase Assay System reagent (Promega) were added. Cells were incubated for 15 min at room temperature on a shaker at 600 rpm. Afterwards, the suspension was mixed by pipetting up and down before 20 μl were transferred into a white 96-well F-bottom plate (Corning). Luminescence was measured using an Infinite F200 PRO plate reader (Tecan) by using an integration time of 100 ms. Luciferase assays were conducted with three technical replicates per sample.

### Flow cytometry

To analyze FAP expression two million R1, R1-FAP or SK-MEL-24 cells were detached with Accutase solution (Sigma-Aldrich), collected in a 15 ml conical tube and spun down at 400 xg for 3 min at 4 °C. The cell pellets were resuspended in 1 ml cold flow buffer (PBS supplemented with 5 % FBS and 0.1 % sodium azide). Next, the cell suspensions were split in two halves and transferred into cold microcentrifuge tubes and washed two more times with cold 500 μl flow buffer. The unstained samples remained in flow buffer, while the stained samples were resuspended in flow buffer supplemented with the primary antibody (anti-FAP human B12, (24) at a dilution of 1:500. Incubation was done for one hour at 4 °C on an end-over-end rotator. Both cell batches, unstained and stained, were washed three times in cold flow buffer and then the secondary antibody (Goat anti-Human IgG H+L Cross-Adsorbed Secondary Antibody, Alexa Fluor™ 488, ThermoFisher Scientific) at a dilution of 1:500 was added. The incubation of the secondary antibody was done for one hour at 4°C on an end-over-end rotator. Cells were washed another three times in cold flow buffer before passed through a 35 μm cell strainer to avoid cell clumps. Flow cytometry was done on a SONY SH800 flow sorter equipped with a 488 nm laser. Gates for the whole cell population and single cell population were adjusted to the different cell types tested. At least 20,000 single cell events were recorded for each sample.

### Statistics

qPCR values were obtained from three independent crude lysate productions. The luciferase data shown were attained by three biological replicates with three technical replicates each. All error bars indicate the standard error of the mean (SEM). For all qPCR and luciferase assay data shown, the differences between DJ and all other capsid variants were tested for statistical significance by one-way ANOVA analysis of variance followed by a Dunnett’s post-hoc test. *p*-values < 0.05 were considered statistically significant (\**p*<0.05; \*\**p* < 0.01; \*\*\**p* < 0.001). All p-values are listed in Table S2-5. Statistical analysis was performed in R (version 4.3.2).

## RESULTS

### Vector design of nanobody insertions into the AAV-DJ capsid for FAP-mediated cell targeting

To enhance the infection efficacy of AAV specifically for FAP receptor-positive cells, but not FAP receptor-negative cells, we inserted a FAP nanobody into the capsid of AAV-DJ (Figure 1A). We chose in total eight different insertion positions on the capsid surface: three positions in the 2-fold valley (Q259, N262, and Q387), two positions in the protruding pore (N328 and N337), two positions in the depression around the pore (N664 and S670), as well as the previously proven insertion position T456 at the tip of the 3-fold protrusion (Figure 1B; (19, 20). Since VP3 makes up the majority of the AAV capsid, we restricted the nanobody embeddings to VP1 or VP2, thereby avoiding potential steric hindrance of capsid formation by too many nanobodies on a single capsid. For insertions into VP1 only, the start codons for VP2 (T138A) and VP3 (M203K, M211L, and M235L) were mutated prior to introducing the FAP nanobody. Further, we generated a rep2-capDJ plasmid with a mutated start codon of VP1 (M1K), complementing VP2 and VP3 expression during AAV production. Likewise, a VP2 only plasmid by mutating start codons of VP1 and VP3 was generated, as well as a plasmid providing only VP1 and VP3 by mutating the VP2 start codon (Figure 1C). For the nanobody insertions, we chose two different types of linkers: one short, symmetric linker pair (SGGGG on both sides) and an asymmetric linker pair, with a long N-terminal 5xSGGGG linker and a short C-terminal GGGGS linker, which was previously used by Eichhoff et al. for their nanobody insertions into VR4 (19). To interrogate AAV re-targeting in the context of adding targeting nanobodies to the AAV capsid alone, we did not mutate AAV-DJ’s heparin binding domain (R587-590), which is often done to attenuate capsid binding to heparan sulfate proteoglycans. Altogether, a total of 32 FAP nanobody variants were generated: 8 positions x 2 different VPs x 2 linker sets.

**Figure 1:**
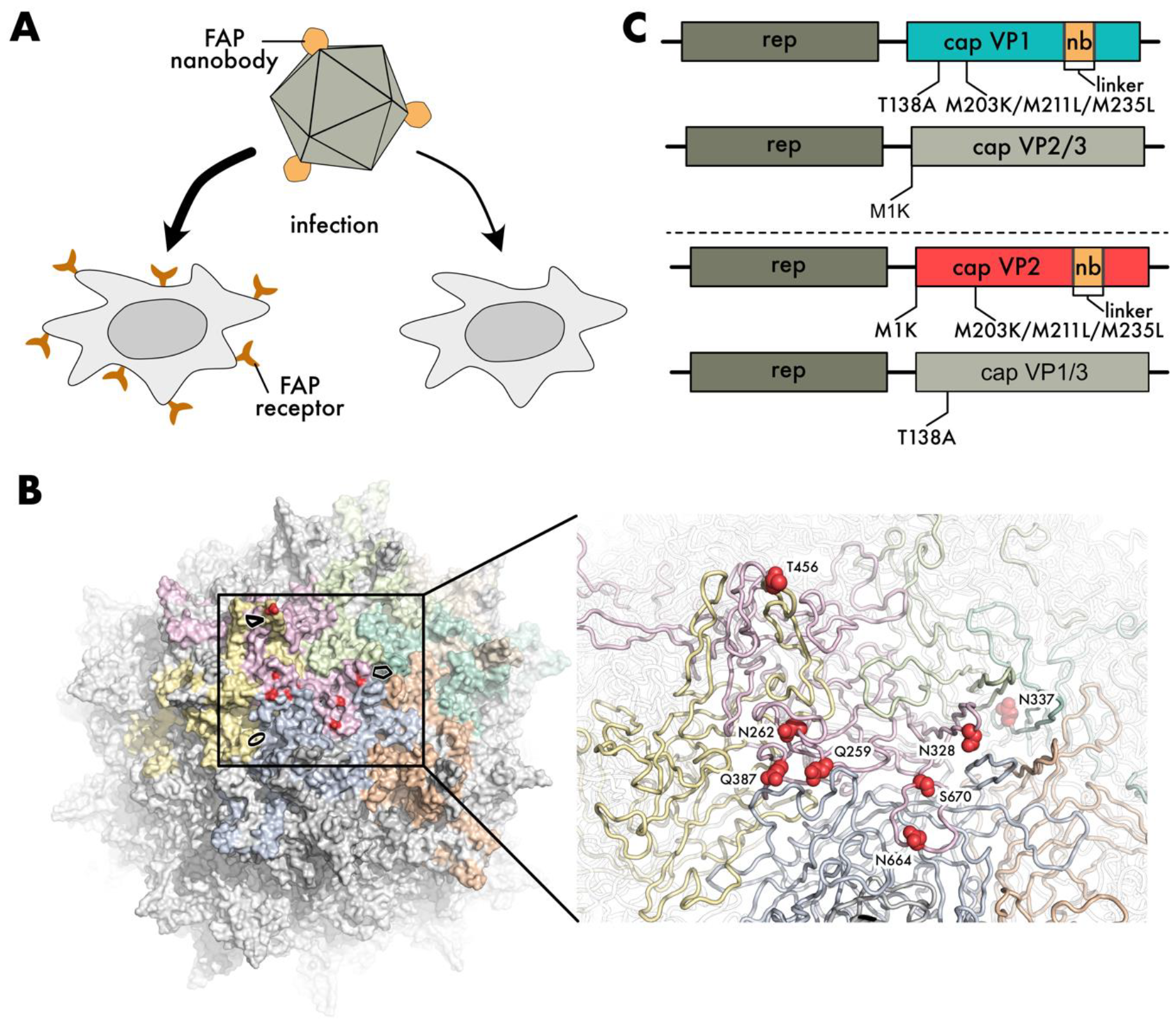
FAP nanobody incorporation at different positions of AAV-DJ. **(A)** Schematic of enhanced infection of FAP receptor-positive cells compared to FAP receptor-negative cells by incorporating a FAP nanobody into the AAV capsid. **(B)** AAV-DJ capsid structure (RCSB PDB 7KFR) from the outside (left) with a zoom-in (right). The eight FAP nanobody insertion positions are highlighted in red. **(C)** Plasmid design of nanobody (nb) insertion into either VP1 (top) or VP2 (bottom) and the respective trans-complementation plasmids for AAV productions.

### FAP nanobody insertions boost transduction in FAP receptor-positive cells

All variants, as well as a DJ control and controls for the split VP expression plasmids (VP1 and VP2/3 or VP2 and VP1/3 expressed from separate plasmids), were produced as crude lysates packaging a CAG promoter-driven NanoLuc payload. Post production, titers were assessed by qPCR. We found that none of the variants had titers statistically significant different to the DJ control. However, we noted a trend that almost invariably all VP1 insertion variants resulted in higher titers than DJ (Figure S2). Next, we used these crude lysates to infect the human prostate cancer cell line CWR-R1-enzalutamide resistant/Fluciferase^+^ (R1) at an MOI of 1x10^3^ vg/cell, which is FAP receptor-negative as verified by flow cytometry (Figure S3A-C). 48 hours post transduction, a luciferase assay using Furimazine substrate for the NanoLuc was conducted and photon counts normalized to the DJ control (Figure 2A). Note, the firefly luciferase (stably expressed by the cell line) and the NanoLuc (delivered as transgene by the AAV) use fully orthogonal substrates, D-luciferin and Furimazine, respectively (30), meaning that the integrated firefly luciferase does not contribute to measured signal. All VP2 insertion variants and nearly all VP1 insertion variants infected R1 cells significantly less efficient than the DJ parent (Figure 2A), indicating that the nanobody insertions to some extend negatively impact the uptake and intracellular processing of the AAV. To test our hypothesis that FAP nanobody insertions into the AAV capsid can boost infection of FAP receptor positive cells, the same AAV samples were used to infected R1-FAP cells, which stably expressed the human FAP receptor (Figure S3D). We found that insertions into positions N262, N328, Q387, T456, N664, and S670 surpassed the infection potency of DJ, independent of the linker type and whether the FAP nanobody insertion was made into VP1 or VP2. The best variant, VP2-N262-asymmetric, infected R1-FAP cells even ∼8-fold better than DJ. Conversely, insertions into positions Q259 and N337 did not enhance infection and showed an equal infection reduction as seen for the assay with the FAP receptor negative R1 cells (Figure 2B). We calculated a FAP nanobody-mediated specificity gain as the ratios of off-to on-target infection efficacy (R1-FAP / R1 infectivity normalized to AAV-DJ). By this metric, our data reveals an up to ∼18-fold improved infection in R1 than in R1-FAP cells for the VP2-N262-asymmetric variant. Even all other variants, that were able to mediate a FAP nanobody-specific infection in R1-FAP cells, showed a specificity gain of >4-fold (Figure 2C). Lowering the dose of transduced AAV to 5x10^2^ and 1x10^2^ vg/cell showed comparable results, demonstrating that the specificity gain is dose independent (Figure S4). Since the R1-FAP cell line is engineered to overexpress FAP, we turned our attention to a cell line with lower endogenous expression (compared to R1-FAP), such as the melanoma cell line SK-MEL-24 (Figure S3E). For the infection of SK-MEL-24 cells, the same trend of infection gain was observed. The symmetric VP1 insertion at position T456 stands out with a 5-fold infection increase, but also VP2 insertions at positions N262, N328, T456, and N664 surpassed the DJ infection potency by at least 2-fold (Figure 2D). To test if the observed specificity gains are FAP nanobody-specific, we conducted an isotype control experiment with a GFP nanobody. To this end, two variants (VP1-T456 and VP1-N664, both with symmetric linkers) were produced and tested in a one-on-one comparison to their FAP nanobody counterparts. As expected, there was no GFP nanobody-specific infection gain as seen for the FAP nanobody (Figure S5).

**Figure 2:**
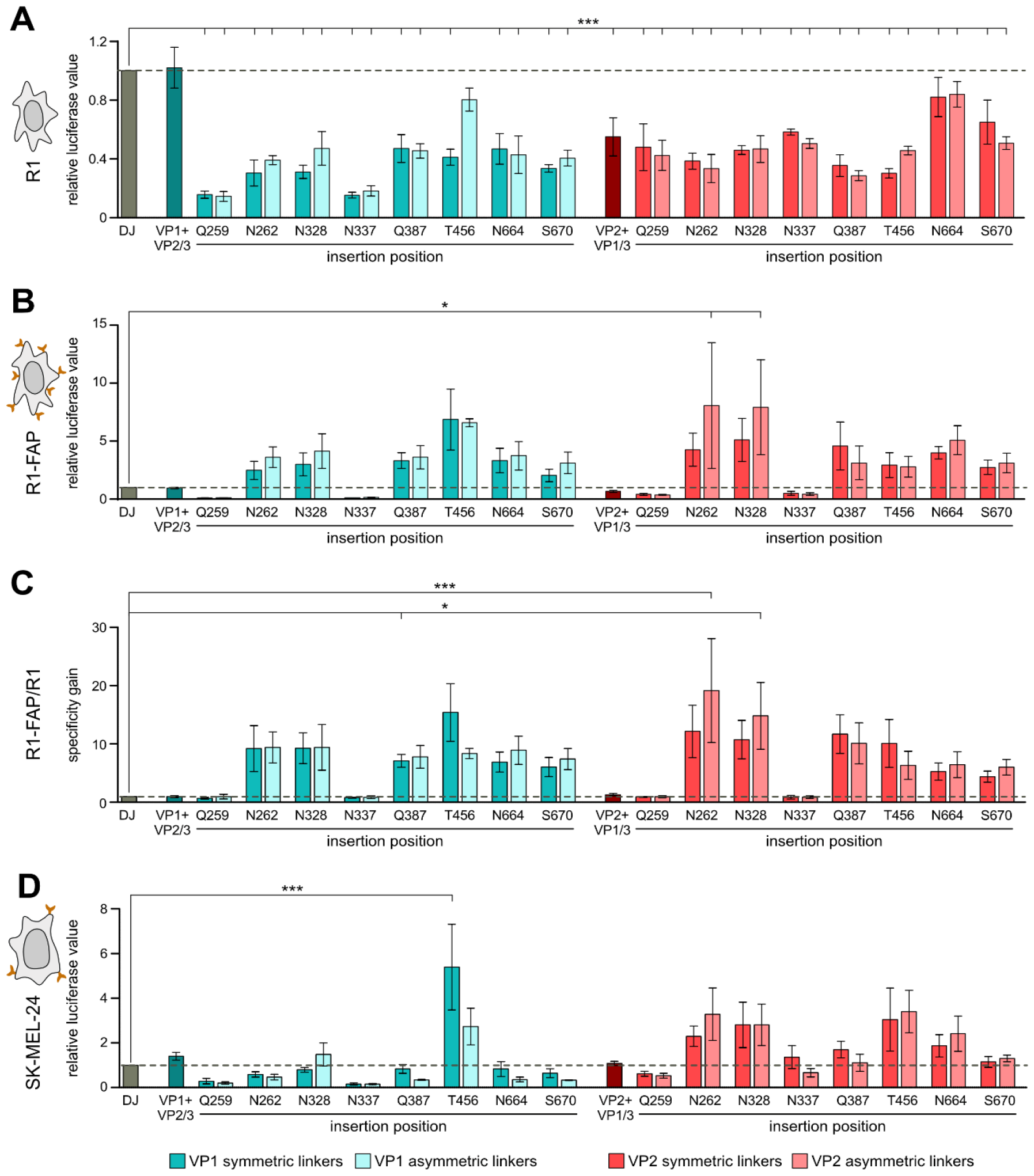
FAP nanobody incorporation at various positions enhances infection of FAP receptor-positive cells. **(A, B, D)** R1 (A), R1-FAP **(B)**, and SK-MEL-24 **(D)** cells were transduced at an MOI of 1x10^3^ vg/cell with the indicated capsid variants, followed by a luciferase assay. Photon counts were normalized to the DJ controls. **(C)** Relative luciferase values from **A** and **B** were divided by each other to calculate the specificity gain. The dashed horizontal line highlights the DJ control level. Data are means ± SEM. \**p* < 0.05, \*\*\**p*<0.001 by one-way ANOVA and a Dunnett’s post hoc test.

### Comparison of infection gain from different cell types reveals the most robust nanobody insertion positions

The differences and similarities between the two different cell lines tested, R1-FAP and SK-MEL-24 are illustrated in Figure 3A. Overall, FAP nanobody embeddings into VP2 yielded a higher infection rate on average than into VP1, except for position T456, the previously published benchmark (19). This position seemed to tolerate nanobody insertions irrespective of the linker. Similarly good as position T456 functioned the variants with insertion at positions N262, N328, and N664, showing the most robust infection gains (Figure 3A). For embeddings into positions Q259 and N337 an infection gain was not observed in either cell line. Mapping of the averaged infection gains for each insertion position into VP1 or VP2 onto the capsid structure revealed no consistent tolerability scheme with regards to the different interfaces of the AAV capsid. For example, all three positions located close to each other within the 2-fold valley (Q259, N262, and Q387) reached very different nanobody-mediated infection gains (Figure 3B).

**Figure 3:**
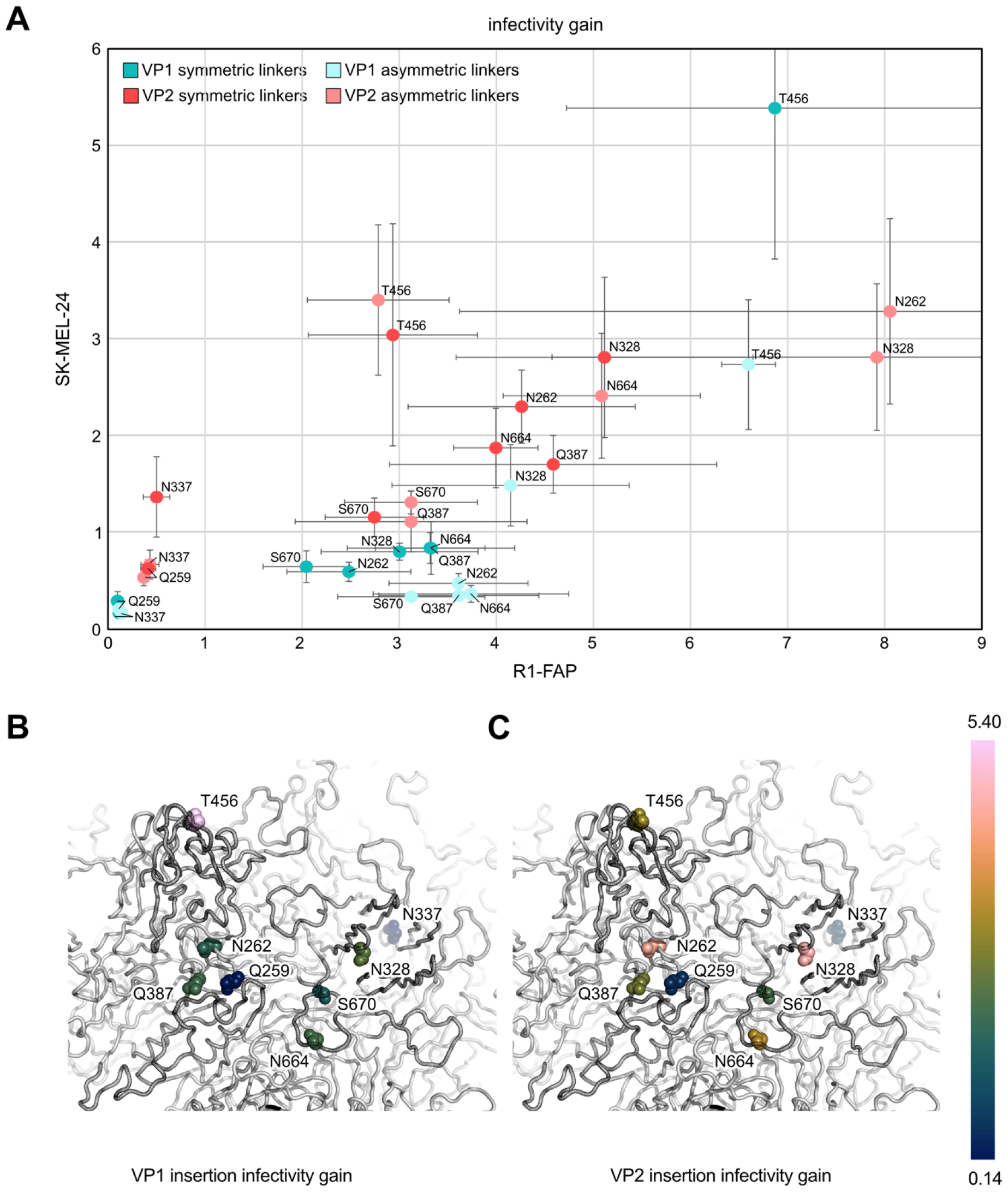
Comparison of infection gain between R1-FAP and SK-MEL-24 cells. **(A)** Scatter plot representing the specificity gain of FAP nanobody insertion variants in R1-FAP cells compared to SK-MEL-24 cells. Data are means ± SEM. **(B, C)** Averaged infectivity gains determined in R1-FAP and SK-MEL-24 from both linker types for FAP-nanobody insertion into VP1 **(B)** and VP2 **(C)** were mapped onto the AAV-DJ capsid structure (RCSB PDB 7KFR).

## DISCUSSION

AAV has been proven to be a suitable vector to efficiently deliver transgenes for cell and gene therapies (1). Nonetheless, broad tissue tropism of natural serotypes hampers its applicability whenever a cell type-specific targeting is of interest. Different capsid engineering approaches have been applied to tackle this issue, including peptide display (9–15), the recovery of AAV ancestors (31), and the insertion of domains with re-targeting abilities (17, 18, 32). In particular, the embedding of nanobodies into the AAV capsid has been strikingly successful in boosting a cell-specific transduction (19, 20).

Nanobodies naturally come with outstanding properties. They are small (15 kDa), stable, have antibody-like binding affinities, and, on top, a low immunogenicity (33). Previous studies had successfully incorporated ARTC2.2, P2X7, CD38, CD4, and GFP nanobodies into VR4 of VP1 and subsequently showed a nanobody-specific uptake by cells expressing the cognate receptor (18–20). Based on our prior domain insertional profiling screen (22), we postulated that nanobodies can be embedded throughout the capsid’s surface, instead of only into the 3-fold protrusion (VR4). Additional options for nanobody insertions may potentially synergize existing VR4 engineering approaches.

In this study, we incorporated a FAP nanobody into eight different positions, covering different surface areas of AAV-DJ. The benchmark position T456, which is part of VR4 was included as a reference. As hypothesized, we could show that AAV capsids can harbor nanobody insertions at various locations, including the 2-fold valley and the protruding 5-fold pore. None of the tested variants showed a reduction in production titer (Figure S2) and six out of the eight chosen positions were able to mediate a FAP receptor-specific infection boost of up to ∼18-fold (Figure 2). The two different linker pairs tested had very minor influence on whether an insertion variant boosted infection or not (Figure 2), which is surprising regarding the fact that the two termini of a nanobody are ∼40 Ångstrom apart and the short symmetric linker could cause sheering forces to the capsid. The choice of engineered VP protein made a bigger difference. We observed increased production titers for almost all VP1 variants, whereas VP2 insertions produced similar to the DJ parent (Figure S2). Despite higher production titers, nanobody incorporations into VP1 had a lower transduction efficacy compared to VP2 insertions (Figure 2,3). We speculate that this effect is caused by a reduced incorporation rate of VP1-nanobody monomers. It has been shown that VP1 is indispensable for the infection process. The unique N-terminus, also known as VP1u, lays inside the capsid and comprises a phospholipase domain (PLA2) as well as a nuclear localization signal. Both play a critical role in endosomal escape and nuclear entry, respectively (34–37). On the one hand, if the capsid contains fewer VP1 monomers, fewer VP1u overhangs occupy the inner cavity of the capsid, possibly facilitating the more efficient packaging of the DNA cargo. On the other hand, fewer VP1 monomers also result in a reduced infection potency. Consequently, the VP2 monomer, for which no crucial functions are known, appears to be the better choice for nanobody insertions. While Eichoff et al. only tested a single insertion into VP1, Hamann et al. also tested N-terminal fusions of a nanobody to VP2, which was less effective than the known VR4 insertion position of VP1 (19, 20). Supporting the notion that engineering VP2 is the more promising route, a previous study by us also tested VP2 insertions into VR4 and found that this insertion was superior compared to the VP1 equivalent (18).

Overall, our FAP nanobody insertion into six out of the eight positions (N262, N328, Q387, T456, N664, and S670) resulted in an infection boost in FAP receptor-positive cells (Figure 2). All these positions have in common that they are on the surface of the capsid, but they are located at very different regions, i.e. at the 5-fold pore (N328), 3-fold protrusion (T456), 2-fold valley (N262 and Q387), and valley surrounding the pore (N664 and S670; Figure 3B,C). One could argue that four of these are part of known VRs and therefore more likely to tolerate insertions (N262 is part of VR1, N328 is part of VR2, Q387 is part of VR3, and T456 is part of VR4), but N664 and S670 also showed a receptor-specific infection and these two positions are neither part of a VR nor a general protrusion (8). Further, three of the position amenable to nanobody insertions (N262, Q387, T456) are known to play a role in the binding of the ubiquitously used AAVR receptor (38, 39). We speculate that insertions right behind these residues are likely to block the AAVR binding and instead promote the FAP receptor binding. Another important receptor used by AAV is heparan sulfate proteoglycan, which is bound by the heparin-binding domain (HBD, R587-R590) of the capsid. Previous studies had removed his binding domain prior to inserting the nanobody to achieve a null-tropism capsid parent to which the infection potency was compared (18–20). We could show here, that even with leaving this known receptor binding site in place, a nanobody-mediated infection boost occurred. For future studies, it could be interesting to investigate in how far deletions or mutations of known receptor binding sites, such as HBD and AAVR, can influence the specificity gain.

Two insertion positions that we tested, N337 and Q259, did not result in any infection gain, although their packaging was not impaired (Figure S2). N337 is partly hidden within the pore and an insertion there could potentially block the externalization of VP1u through the pore during infection. Q259 lays within the 2-fold valley closely abutting a neighboring VP subunit (unlike the nearby N262). Prior work has shown that interfacial dynamics at the twofold axis play a role in externalization of VP1u during infection, which may explain the lack of infectivity of insertion variant at this position (40).

The FAP nanobody used in this study can be used to direct AAV to cancer or cancer-associated cells with a high FAP expression profile (23), but further experiments *in vivo* are required for validation. Notably, the straightforward replacement of the GFP nanobody in our previous study (22) with a FAP nanobody here suggests that nanobody swapping at the 2-fold valley and 5-fold axis hotspots is a viable strategy to rapidly diversify AAV tropism. With more and more nanobodies being developed, the here demonstrated tolerability of nanobody insertions at various locations within the AAV capsid can further expand the applicability of nanobody-directed cell-specific targeting.

## DATA AVAILABILITY

All plasmids are available upon request. Flow cytometry data is available on FlowRepository (Repository ID: FR-FCM-Z789).

## SUPPLEMENTARY DATA

Supplementary Data are available at NAR Molecular Medicine online.

## AUTHOR CONTRIBUTIONS

MDH and DS conceived the study with input from JPG and AL. MDH conducted experiments. MDH and DS analyzed the data and authored the manuscript. All authors have given approval to the final version of the manuscript.

