## Supplementary Data for "Unlocking Precision Gene Therapy: Harnessing AAV Tropism with Nanobody Swapping at Capsid Hotspots"

FAP nanobody with asymmetric linkers

GGGGSGGGGSGGGGSGGGGSGGGGSQVQLVESGGGLVQPGGSLRLSCAASGF
TFNNYWVYWARQAPGKGLEWVSSMNIADGTTYYAASVKGRFTISSDNAKNTVYLQ
MNSLKSEDTAVYYCAKGVVGPVGNGGLYLGKGTLVTVSGGGGA

FAP nanobody with symmetric linkers

SGGGGQVQLVESGGGLVQPGGSLRLSCAASGFTFNNYWVYWARQAPGKGLEWV
SSMNIADGTTYYAASVKGRFTISSDNAKNTVYLQMNSLKSEDTAVYYCAKGVVGPV
GNGGLYLGKGTLVTVSGGGGS

GFP nanobody with symmetric linkers

SGGGGQVQLVESGGALVQPGGSLRLSCAASGFPVNRYSMRWYRQAPGKEREWV
AGMSSAGDRSSYEDSVKGRFTISRDDARNTVYLQMNSLKPEDTAVYYCNVNVGFE
YWGQGTQVTVSSKGGGGS

**Figure S1**

Protein sequences of FAP (yellow) and GFP (purple) nanobodies with linkers (grey).

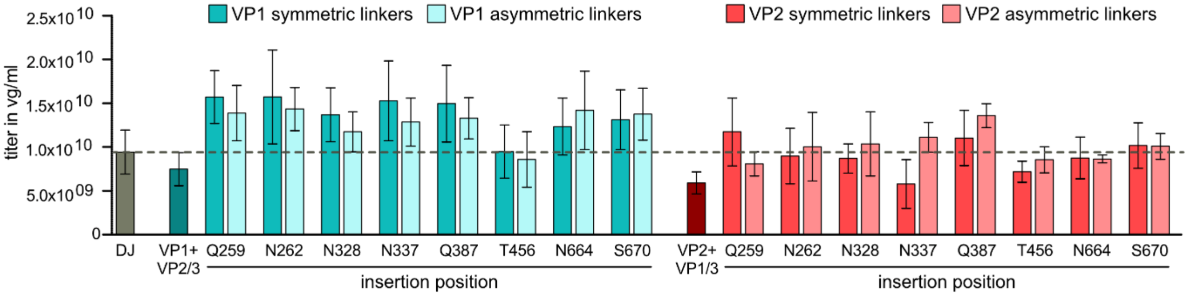


**Figure S2**

Production titers of nanobody insertion variants. Quantification of AAV crude lysate titers by qPCR. The dashed horizontal line highlights the DJ control level. Data are means ± SEM. Differences are not significant for any capsid variant compared to DJ by one-way ANOVA and a Dunnett's post hoc test.

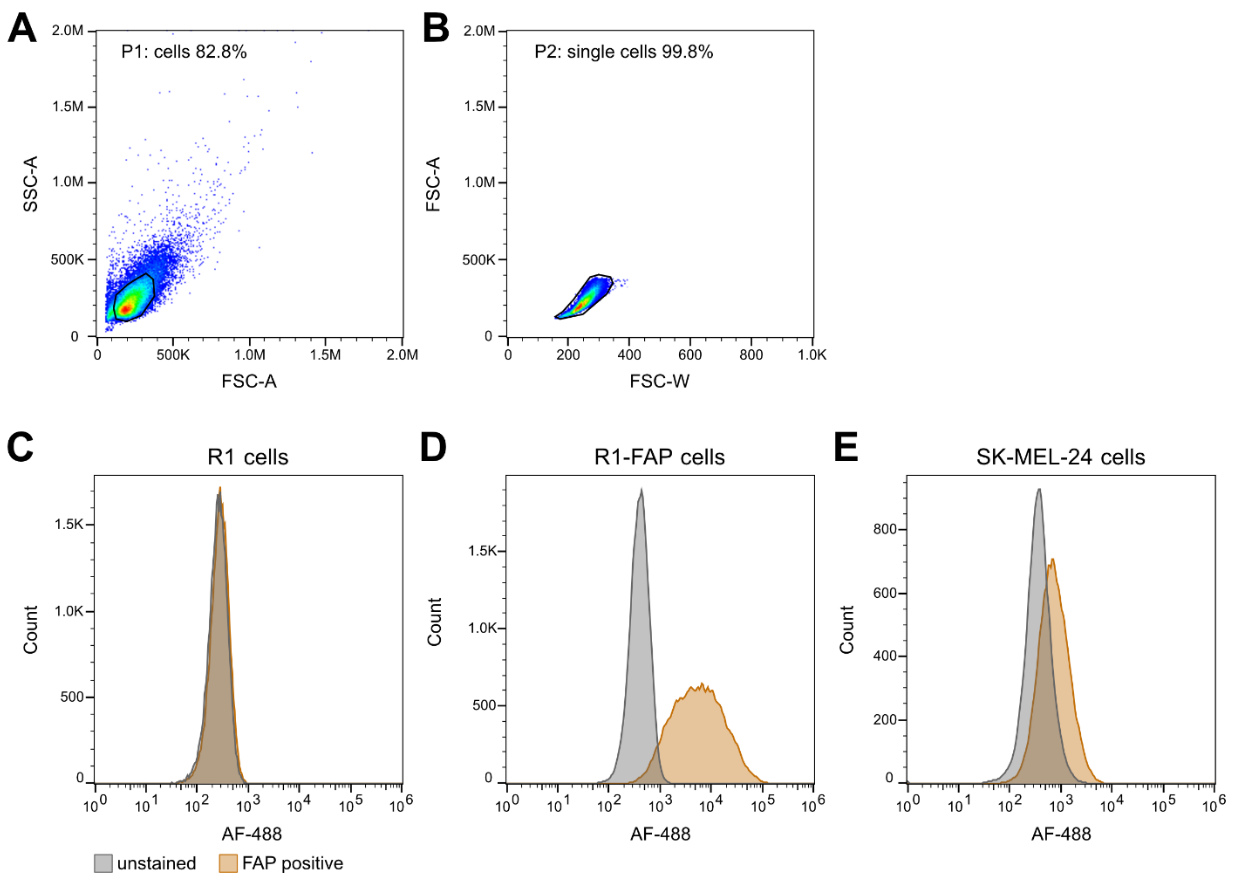


**Figure S3**

FAP receptor expression profiles. (A) Whole cells (gate P1) were gated on forward scattering area (FSC-A) and side scattering area (SSC-A). (B) Forward scattering width (FSC-W) and forward scattering area (FSC-A) were used to gate single cells (P2). (C-E) Histograms of unstained or FAP receptor stained single cell populations of R1 cells (C), R1-FAP cells (D), and SK-MEL-24 cells (E).

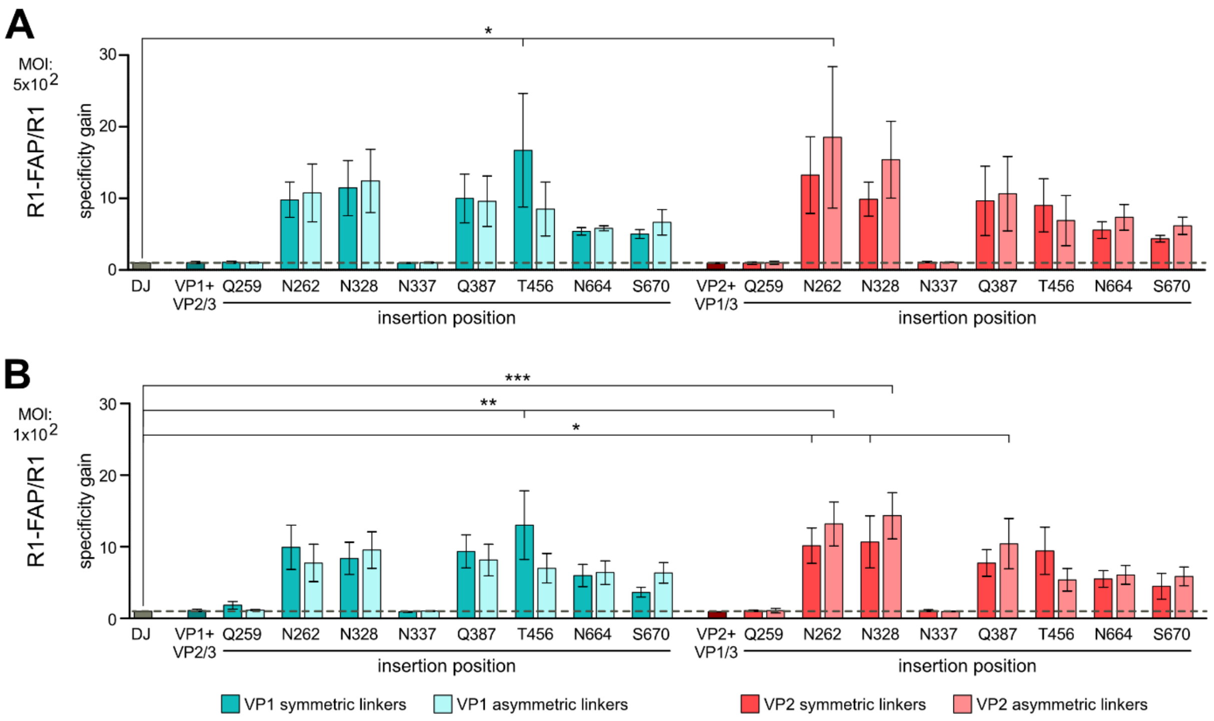


**Figure S4**

Specificity gain is dose-independent. (A-B) R1 and R1-FAP cells were transduced at an MOI of $5x10^2$ (A) or $1x10^2$ (B) vg/cell with the indicated capsid variants, followed by a luciferase assay. First, photon counts were normalized to the DJ controls of each cell line and second the R1-FAP values were divided by the R1 values to calculate the specificity gain. The dashed horizontal line highlights the DJ control level. Data are means ± SEM. *$p < 0.05$, **$p < 0.01$, ***$p<0.001$ by one-way ANOVA and a Dunnett's post-hoc test.

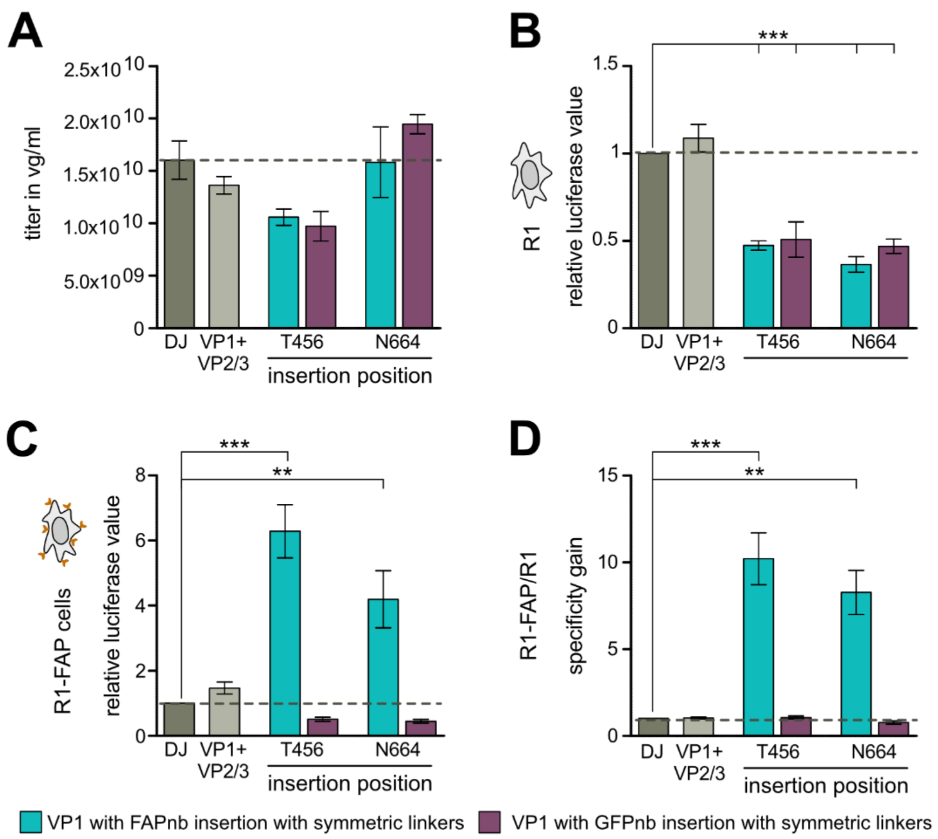


**Figure S5**

Specificity gain of infection is caused specifically by the FAP nanobody incorporated into the capsid. (A) Quantification of AAV crude lysate titers by qPCR. (B-C) R1 cells (B) and R1-FAP cells (C) were transduced at an MOI of $1x10^3$ vg/cell with the indicated capsid variants, followed by a luciferase assay. Photon counts were normalized to the DJ controls. (D) Relative luciferase values from B and C were divided by each other to calculate the specificity gain. The dashed horizontal line highlights the DJ control level Data are means ± SEM. ***p* < 0.01, ****p*<0.001 by one-way ANOVA and a Dunnett's post-hoc test.

### Table S1

Plasmids used in this study. sym, symmetric. asym, asymmetric. nb, nanobody.

| # | Name | Encodes/Use | Origin |
|---|---|---|---|
| 1 | nanoLuc | nanoLuciferase packaging | Schmidt lab |
| 2 | Adeno-helper | Adenohelper genes for AAV productions | Cell Biolabs |
| 3 | AAV-DJ | rep2-capDJ for packaging of AAV | Cell Biolabs |
| 4 | DJ-VP1 | rep2-capDJ with mutated start codons for VP2/VP3 | Schmidt lab (MDH) |
| 5 | DJ-VP2 | rep2-capDJ with mutated start codons for VP1/VP3 | Schmidt lab (MDH) |
| 6 | DJ-VP2/3 | rep2-capDJ with mutated start codon for VP1 | Schmidt lab (MDH) |
| 7 | DJ-VP1/3 | rep2-capDJ with mutated start codon for VP2 | Schmidt lab (MDH) |
| 8 | DJ-VP1-T456-FAPnb-sym | capDJ-VP1 with symmetric linker FAP-nanobody insertion at T456 | Schmidt lab (MDH) |
| 9 | DJ-VP1-T456-FAPnb-asym | capDJ-VP1 with asymmetric linker FAP-nanobody insertion at T456 | Schmidt lab (MDH) |
| 10 | DJ-VP1-N664-FAPnb-sym | capDJ-VP1 with symmetric linker FAP-nanobody insertion at N664 | Schmidt lab (MDH) |
| 11 | DJ-VP1-N664-FAPnb-asym | capDJ-VP1 with asymmetric linker FAP-nanobody insertion at N664 | Schmidt lab (MDH) |
| 12 | DJ-VP1-S670-FAPnb-sym | capDJ-VP1 with symmetric linker FAP-nanobody insertion at S670 | Schmidt lab (MDH) |
| 13 | DJ-VP1-S670-FAPnb-asym | capDJ-VP1 with asymmetric linker FAP-nanobody insertion at S670 | Schmidt lab (MDH) |
| 14 | DJ-VP1-N337-FAPnb-sym | capDJ-VP1 with symmetric linker FAP-nanobody insertion at N337 | Schmidt lab (MDH) |
| 15 | DJ-VP1-N337-FAPnb-asym | capDJ-VP1 with asymmetric linker FAP-nanobody insertion at N337 | Schmidt lab (MDH) |
| 16 | DJ-VP1-N328-FAPnb-sym | capDJ-VP1 with symmetric linker FAP-nanobody insertion at N328 | Schmidt lab (MDH) |
| 17 | DJ-VP1-N328-FAPnb-asym | capDJ-VP1 with asymmetric linker FAP-nanobody insertion at N328 | Schmidt lab (MDH) |
| 18 | DJ-VP1-Q259-FAPnb-sym | capDJ-VP1 with symmetric linker FAP-nanobody insertion at Q259 | Schmidt lab (MDH) |
| 19 | DJ-VP1-Q259-FAPnb-asym | capDJ-VP1 with asymmetric linker FAP-nanobody insertion at Q259 | Schmidt lab (MDH) |
| 20 | DJ-VP1-N262-FAPnb-sym | capDJ-VP1 with symmetric linker FAP-nanobody insertion at N262 | Schmidt lab (MDH) |
| 21 | DJ-VP1-N262-FAPnb-asym | capDJ-VP1 with asymmetric linker FAP-nanobody insertion at N262 | Schmidt lab (MDH) |
| 22 | DJ-VP1-Q387-FAPnb-sym | capDJ-VP1 with symmetric linker FAP-nanobody insertion at Q387 | Schmidt lab (MDH) |
| 23 | DJ-VP1-Q387-FAPnb-asym | capDJ-VP1 with asymmetric linker FAP-nanobody insertion at Q387 | Schmidt lab (MDH) |
| *24* | DJ-VP2-T456-FAPnb-sym | capDJ-VP2 with symmetric linker FAP-nanobody insertion at T456 | Schmidt lab (MDH) |
| *25* | DJ-VP2-T456-FAPnb-asym | capDJ-VP2 with asymmetric linker FAP-nanobody insertion at T456 | Schmidt lab (MDH) |
| *26* | DJ-VP2-N664-FAPnb-sym | capDJ-VP2 with symmetric linker FAP-nanobody insertion at N664 | Schmidt lab (MDH) |
| *27* | DJ-VP2-N664-FAPnb-asym | capDJ-VP2 with asymmetric linker FAP-nanobody insertion at N664 | Schmidt lab (MDH) |
| *28* | DJ-VP2-S670-FAPnb-sym | capDJ-VP2 with symmetric linker FAP-nanobody insertion at S670 | Schmidt lab (MDH) |
| *29* | DJ-VP2-S670-FAPnb-asym | capDJ-VP2 with asymmetric linker FAP-nanobody insertion at S670 | Schmidt lab (MDH) |
| *30* | DJ-VP2-N337-FAPnb-sym | capDJ-VP2 with symmetric linker FAP-nanobody insertion at N337 | Schmidt lab (MDH) |
| *31* | DJ-VP2-N337-FAPnb-asym | capDJ-VP2 with asymmetric linker FAP-nanobody insertion at N337 | Schmidt lab (MDH) |
| *32* | DJ-VP2-N328-FAPnb-sym | capDJ-VP2 with symmetric linker FAP-nanobody insertion at N328 | Schmidt lab (MDH) |
| *33* | DJ-VP2-N328-FAPnb-asym | capDJ-VP2 with asymmetric linker FAP-nanobody insertion at N328 | Schmidt lab (MDH) |
| *34* | DJ-VP2-Q259-FAPnb-sym | capDJ-VP2 with symmetric linker FAP-nanobody insertion at Q259 | Schmidt lab (MDH) |
| *35* | DJ-VP2-Q259-FAPnb-asym | capDJ-VP2 with asymmetric linker FAP-nanobody insertion at Q259 | Schmidt lab (MDH) |
| *36* | DJ-VP2-N262-FAPnb-sym | capDJ-VP2 with symmetric linker FAP-nanobody insertion at N262 | Schmidt lab (MDH) |
| *37* | DJ-VP2-N262-FAPnb-asym | capDJ-VP2 with asymmetric linker FAP-nanobody insertion at N262 | Schmidt lab (MDH) |
| *38* | DJ-VP2-Q387-FAPnb-sym | capDJ-VP2 with symmetric linker FAP-nanobody insertion at Q387 | Schmidt lab (MDH) |
| *39* | DJ-VP2-Q387-FAPnb-asym | capDJ-VP2 with asymmetric linker FAP-nanobody insertion at Q387 | Schmidt lab (MDH) |
| 40 | DJ-VP1-T456-GFPnb-sym | capDJ-VP1 with symmetric linker GFP-nanobody insertion at T456 | Schmidt lab (MDH) |
| 41 | DJ-VP1-N664-GFPnb-sym | capDJ-VP1 with symmetric linker GFP-nanobody insertion at N664 | Schmidt lab (MDH) |

**Table S2**

p-values of one-way ANOVA with Dunnett's post-hoc test from data in Figure 2A-D (n.s.: not significant).

| | | | Figure 2A | | Figure 2B | | Figure 2C | | Figure 2D | |
|---|---|---|---|---|---|---|---|---|---|---|
| variant | | | p value | significance | p value | significance | p value | significance | p value | significance |
| VP1 + VP2/3 | | | 1.0000 | n.s. | 1.0000 | n.s. | 1.0000 | n.s. | 1.0000 | n.s. |
| VP1 insertion | Q259 | symmetric | 0.8691 | n.s. | 0.0000 | *** | 1.0000 | n.s. | 1.0000 | n.s. |
| | | asymmetric | 0.9956 | n.s. | 0.0000 | *** | 1.0000 | n.s. | 1.0000 | n.s. |
| | N262 | symmetric | 0.8653 | n.s. | 0.0000 | *** | 1.0000 | n.s. | 0.5008 | n.s. |
| | | asymmetric | 0.9855 | n.s. | 0.0000 | *** | 0.9806 | n.s. | 0.4646 | n.s. |
| | N328 | symmetric | 0.9978 | n.s. | 0.0000 | *** | 0.9995 | n.s. | 0.4891 | n.s. |
| | | asymmetric | 1.0000 | n.s. | 0.0004 | *** | 0.8950 | n.s. | 0.4662 | n.s. |
| | N337 | symmetric | 0.9199 | n.s. | 0.0000 | *** | 1.0000 | n.s. | 1.0000 | n.s. |
| | | asymmetric | 0.9999 | n.s. | 0.0000 | *** | 1.0000 | n.s. | 1.0000 | n.s. |
| | Q387 | symmetric | 0.9498 | n.s. | 0.0005 | *** | 0.9954 | n.s. | 0.0163 | * |
| | | asymmetric | 0.9995 | n.s. | 0.0003 | *** | 0.9805 | n.s. | 0.6596 | n.s. |
| | T456 | symmetric | 1.0000 | n.s. | 0.7346 | n.s. | 0.1281 | n.s. | 0.9064 | n.s. |
| | | asymmetric | 1.0000 | n.s. | 0.0000 | *** | 0.1689 | n.s. | 0.5560 | n.s. |
| | N664 | symmetric | 0.9899 | n.s. | 0.0004 | *** | 0.9952 | n.s. | 0.9762 | n.s. |
| | | asymmetric | 1.0000 | n.s. | 0.0001 | *** | 0.9681 | n.s. | 0.8324 | n.s. |
| | S670 | symmetric | 0.9998 | n.s. | 0.0000 | *** | 1.0000 | n.s. | 0.8782 | n.s. |
| | | asymmetric | 0.9972 | n.s. | 0.0000 | *** | 0.9987 | n.s. | 0.7695 | n.s. |
| VP2 + VP1/3 | | | 0.9999 | n.s. | 1.0000 | n.s. | 1.0000 | n.s. | 1.0000 | n.s. |
| VP2 insertion | Q259 | symmetric | 1.0000 | n.s. | 0.0007 | *** | 1.0000 | n.s. | 1.0000 | n.s. |
| | | asymmetric | 1.0000 | n.s. | 0.0001 | *** | 1.0000 | n.s. | 1.0000 | n.s. |
| | N262 | symmetric | 1.0000 | n.s. | 0.0000 | *** | 0.8656 | n.s. | 0.1285 | n.s. |
| | | asymmetric | 1.0000 | n.s. | 0.0000 | *** | 0.0321 | * | 0.0009 | *** |
| | N328 | symmetric | 1.0000 | n.s. | 0.0004 | *** | 0.5736 | n.s. | 0.2635 | n.s. |
| | | asymmetric | 1.0000 | n.s. | 0.0005 | *** | 0.0384 | * | 0.0247 | * |
| | N337 | symmetric | 1.0000 | n.s. | 0.0128 | * | 1.0000 | n.s. | 1.0000 | n.s. |
| | | asymmetric | 1.0000 | n.s. | 0.0015 | ** | 1.0000 | n.s. | 1.0000 | n.s. |
| | Q387 | symmetric | 1.0000 | n.s. | 0.0000 | *** | 0.7631 | n.s. | 0.1655 | n.s. |
| | | asymmetric | 0.9982 | n.s. | 0.0000 | *** | 0.9987 | n.s. | 0.3508 | n.s. |
| | T456 | symmetric | 1.0000 | n.s. | 0.0000 | *** | 0.9997 | n.s. | 0.3527 | n.s. |
| | | asymmetric | 1.0000 | n.s. | 0.0003 | *** | 0.9990 | n.s. | 0.9594 | n.s. |
| | N664 | symmetric | 1.0000 | n.s. | 0.8444 | n.s. | 0.9284 | n.s. | 0.9971 | n.s. |
| | | asymmetric | 1.0000 | n.s. | 0.9284 | n.s. | 0.5832 | n.s. | 0.9498 | n.s. |
| | S670 | symmetric | 1.0000 | n.s. | 0.0629 | n.s. | 1.0000 | n.s. | 0.9999 | n.s. |
| | | asymmetric | 1.0000 | n.s. | 0.0015 | ** | 0.9987 | n.s. | 0.9782 | n.s. |

**Table S3**

p-values of one-way ANOVA with Dunnett's post-hoc test from data in Figure 3A (n.s.: not significant).

| variant | | | p value | significance |
|---|---|---|---|---|
| VP1 + VP2/3 | | | 1.0000 | n.s. |
| VP1 insertion | Q259 | symmetric | 0.9999 | n.s. |
| | | asymmetric | 0.9995 | n.s. |
| | N262 | symmetric | 1.0000 | n.s. |
| | | asymmetric | 1.0000 | n.s. |
| | N328 | symmetric | 1.0000 | n.s. |
| | | asymmetric | 1.0000 | n.s. |
| | N337 | symmetric | 0.9990 | n.s. |
| | | asymmetric | 0.9989 | n.s. |
| | Q387 | symmetric | 1.0000 | n.s. |
| | | asymmetric | 1.0000 | n.s. |
| | T456 | symmetric | 0.0000 | *** |
| | | asymmetric | 0.4992 | n.s. |
| | N664 | symmetric | 1.0000 | n.s. |
| | | asymmetric | 1.0000 | n.s. |
| | S670 | symmetric | 1.0000 | n.s. |
| | | asymmetric | 1.0000 | n.s. |
| VP2 + VP1/3 | | | 1.0000 | n.s. |
| VP2 insertion | Q259 | symmetric | 1.0000 | n.s. |
| | | asymmetric | 1.0000 | n.s. |
| | N262 | symmetric | 0.8728 | n.s. |
| | | asymmetric | 0.1533 | n.s. |
| | N328 | symmetric | 0.4362 | n.s. |
| | | asymmetric | 0.4365 | n.s. |
| | N337 | symmetric | 1.0000 | n.s. |
| | | asymmetric | 1.0000 | n.s. |
| | Q387 | symmetric | 1.0000 | n.s. |
| | | asymmetric | 1.0000 | n.s. |
| | T456 | symmetric | 0.2722 | n.s. |
| | | asymmetric | 0.1133 | n.s. |
| | N664 | symmetric | 0.9982 | n.s. |
| | | asymmetric | 0.7871 | n.s. |
| | S670 | symmetric | 1.0000 | n.s. |
| | | asymmetric | 1.0000 | n.s. |

**Table S4**

p-values of one-way ANOVA with Dunnett's post-hoc test from data in Figure S2A-B (n.s.: not significant).

| | | | Figure S2A | | Figure S2B | |
|---|---|---|---|---|---|---|
| variant | | | p value | significance | p value | significance |
| VP1 + VP2/3 | | | 1.0000 | n.s. | 1.0000 | n.s. |
| VP1 insertion | Q259 | symmetric | 1.0000 | n.s. | 1.0000 | n.s. |
| | | asymmetric | 1.0000 | n.s. | 1.0000 | n.s. |
| | N262 | symmetric | 0.6466 | n.s. | 0.0616 | n.s. |
| | | asymmetric | 0.4952 | n.s. | 0.3122 | n.s. |
| | N328 | symmetric | 0.3855 | n.s. | 0.2038 | n.s. |
| | | asymmetric | 0.2756 | n.s. | 0.0844 | n.s. |
| | N337 | symmetric | 1.0000 | n.s. | 1.0000 | n.s. |
| | | asymmetric | 1.0000 | n.s. | 1.0000 | n.s. |
| | Q387 | symmetric | 0.6176 | n.s. | 0.0986 | n.s. |
| | | asymmetric | 0.6800 | n.s. | 0.2395 | n.s. |
| | T456 | symmetric | 0.0359 | * | 0.0028 | ** |
| | | asymmetric | 0.8426 | n.s. | 0.4740 | n.s. |
| | N664 | symmetric | 0.9997 | n.s. | 0.6353 | n.s. |
| | | asymmetric | 0.9986 | n.s. | 0.7422 | n.s. |
| | S670 | symmetric | 0.9999 | n.s. | 0.9997 | n.s. |
| | | asymmetric | 0.9875 | n.s. | 0.6401 | n.s. |
| VP2 + VP1/3 | | | 1.0000 | n.s. | 1.0000 | n.s. |
| VP2 insertion | Q259 | symmetric | 1.0000 | n.s. | 1.0000 | n.s. |
| | | asymmetric | 1.0000 | n.s. | 1.0000 | n.s. |
| | N262 | symmetric | 0.1966 | n.s. | 0.0498 | * |
| | | asymmetric | 0.0123 | * | 0.0023 | ** |
| | N328 | symmetric | 0.6349 | n.s. | 0.0315 | * |
| | | asymmetric | 0.0727 | n.s. | 0.0006 | *** |
| | N337 | symmetric | 1.0000 | n.s. | 1.0000 | n.s. |
| | | asymmetric | 1.0000 | n.s. | 1.0000 | n.s. |
| | Q387 | symmetric | 0.6731 | n.s. | 0.3121 | n.s. |
| | | asymmetric | 0.5124 | n.s. | 0.0392 | * |
| | T456 | symmetric | 0.7717 | n.s. | 0.0933 | n.s. |
| | | asymmetric | 0.9798 | n.s. | 0.8785 | n.s. |
| | N664 | symmetric | 0.9994 | n.s. | 0.8519 | n.s. |
| | | asymmetric | 0.9579 | n.s. | 0.7170 | n.s. |
| | S670 | symmetric | 1.0000 | n.s. | 0.9857 | n.s. |
| | | asymmetric | 0.9960 | n.s. | 0.7751 | n.s. |

**Table S5**

p-values of one-way ANOVA with Dunnett's post-hoc test from data in Figure 3A-D (n.s.: not significant).

| | | | Figure S3A | | Figure S3B | | Figure S3C | | Figure S3D | |
|---|---|---|---|---|---|---|---|---|---|---|
| variant | | | p value | significance | p value | significance | p value | significance | p value | significance |
| VP1 + VP2/3 | | | 0.8032 | n.s. | 0.7352 | n.s. | 0.9409 | n.s. | 1.0000 | n.s. |
| VP1 insertion | FAPnb | T456 | 0.1714 | n.s. | 0.0002 | *** | 0.0000 | *** | 0.0000 | *** |
| | | N664 | 1 | n.s. | 0.0000 | *** | 0.0029 | ** | 0.0001 | ** |
| | GFPnb | T456 | 0.0985 | n.s. | 0.0003 | *** | 0.929 | n.s. | 1.0000 | n.s. |
| | | N664 | 0.5395 | n.s. | 0.0001 | *** | 0.8907 | n.s. | 0.9997 | n.s. |
